# Shared Transcriptomic Signatures in Perilesional and Contralesional Cortex

**DOI:** 10.64898/2026.02.02.703306

**Authors:** Dene Betz, Victoria A Alers, Matthew Kenwood, Kielen R Zuurbier, Rebeca Coimbra, Priscilla Rhoton, Erik J Plautz, Peter M Douglas, Denise MO Ramirez, Ann M Stowe, Mark P Goldberg

## Abstract

Stroke induces a transient period of heightened plasticity during which functional recovery is most pronounced. Work in rodent models of stroke has identified key mechanisms in both the ipsilesional and contralesional cortex that contribute to functional and structural post-stroke plasticity. To date, most gene expression studies have focused on the infarct core and the immediately surrounding tissue, peri-lesional cortex (PLC). We sought to understand whether the contralesional cortex (CLC), a region that shows robust structural and molecular remodeling despite its distance from the lesion, mounts a distinct transcriptional response to stroke. Direct comparisons of molecular pathways governing these regions, particularly across sexes, are limited. To address this gap, we performed bulk RNA sequencing of the PLC and CLC at 7 days post-stroke, a critical time point for initiating repair, in male and female mice. Our results indicate that despite distinct positions from the lesion, both regions share a robust upregulation of inflammatory signaling, with Gene Ontology enrichment indicating activation of cytokine signaling, leukocyte activation, and gliogenesis pathways. Further analysis of this shared gene expression signature revealed reactive microglia signaling as the dominant pathway. Surprisingly, the CLC did not show a distinct transcriptional response. These findings were consistent across males and females, which also showed similar CLC-derived corticospinal tract axonal sprouting at 6 weeks post-stroke. Together, these findings support a shared microglia-centered neuroinflammatory transcriptional response in the PLC and CLC and suggest that microglial reactivity is a key early process for post-stroke cortical plasticity in both male and female mice.

## Background

Stroke patients present with a wide range of deficits, often followed by a remarkable period of improvement that spans the first 3-6 months after stroke, known as the critical window (Bernhardt et al., 2017; Grefkes & Fink, 2020). During this time, the brain exhibits a heightened capacity for neuronal plasticity that supports functional reorganization. However, most individuals experience a plateau in recovery and are left with chronic deficits that reduce independence and quality of life (Donkor, 2018; Feigin et al., 2019). Investigations into the mechanisms underlying this critical window may help drive the discovery of new therapies for stroke recovery. Preclinical models of stroke have been instrumental in advancing our understanding of these mechanisms. Rodent models replicate key aspects of human stroke recovery and are highly amenable to molecular and genetic manipulation, making them ideal for dissecting the biological processes that underlie post-stroke plasticity. A consistent finding across models is that recovery depends heavily on the capacity of spared neural circuits to rewire and compensate for damaged networks, which occurs through numerous forms of plasticity, including dendritic arborization, synaptic tuning, and axonal sprouting (Brown et al., 2009; Dancause et al., 2005; Okabe et al., 2025; Ueno et al., 2012; Zhang et al., 2010). This is especially evident in the context of focal motor cortical strokes, which elicit robust plasticity in both local and remote regions (Carmichael et al., 2001; Kaiser et al., 2019; Poinsatte et al., 2025).

Among regions undergoing post-stroke plasticity, the peri-lesional cortex (PLC) and contralesional cortex (CLC) are the most well characterized. In the PLC, recruitment of the somatosensory and premotor cortex enables the formation of new connections that directly drive recovery (Brown et al., 2009; Li et al., 2015; Li et al., 2010; Overman et al., 2012; Plautz et al., 2023). In contrast, the role of the CLC is more complex (Sato et al., 2021). In patients, altered neuronal activity has been frequently observed and associated with both beneficial and maladaptive outcomes (Buetefisch, 2015). Rodent models support these findings, demonstrating altered gene expression, axonal sprouting, dendritic remodeling, and changes in neuronal excitability within CLC that differentially affect outcomes (Götz et al., 2023; Urban et al., 2012). Some experimental paradigms, particularly those involving large infarcts encompassing motor and sensory cortex, require CLC to drive recovery (Aswendt et al., 2021; Ueno et al., 2012), while others show no relationship or even an inhibition of functional improvement (Bice et al., 2022; Jones & Schallert, 1994). This apparent contradiction may be resolved by more precise characterization of the CLC transcriptome after stroke, as structural and molecular changes in the CLC occur in parallel with those in the PLC, consistent with a stroke-induced molecular program that coordinates plasticity across regions.

Although the PLC and CLC exhibit broadly similar responses to stroke, relatively few studies have directly compared their transcriptomes, highlighting an opportunity to better understand regional plasticity programs (Buga et al., 2008; Filippenkov et al., 2023; Ito et al., 2018; Keyvani et al., 2002). The PLC lies adjacent to the infarct core and glial scar and is subject to drastic shifts in its local microenvironment from evolving inflammatory signaling. These features are thought to create a neurogenic niche that supports structural remodeling (Carmichael et al., 2005; Nudo et al., 1996). In contrast, CLC is anatomically distant from the lesion and does not experience direct ischemic injury or the same degree of damage-related signaling. How the CLC engages a similarly robust and functionally relevant plasticity program under these conditions remains unclear. A direct comparison of the molecular landscape across these two regions is needed to determine whether shared transcriptional programs support recovery or whether distinct region-specific mechanisms drive remodeling.

Sex has been rigorously investigated in stroke epidemiology, acute injury, and early outcomes; however, its contribution to recovery, particularly the molecular mechanisms that coordinate repair, remains incompletely understood. Clinical studies report inconsistent associations, likely reflecting confounding factors such as lifespan differences, social isolation, and comorbidity burden (Poggesi et al., 2021). In contrast, experimental evidence supports a clear biological role for sex in processes implicated in recovery, including immune signaling and blood-brain barrier integrity (Banerjee & McCullough, 2022; Chisholm & Sohrabji, 2016; Dotson et al., 2015; Manwani et al., 2013; Morrison & Filosa, 2016). Despite this, few studies have systematically examined how males and females differ in their transcriptional response to stroke, particularly across brain regions critical for motor recovery. In this study, we aimed to define and compare the transcriptomic landscape of the peri- and contra-lesional cortex at 7 days post-stroke, a time point corresponding to the initiation of post-stroke plasticity. We further compared these molecular programs across sexes and evaluated the functional relevance of these molecular changes by quantifying axonal sprouting in both male and female mice.

## Materials and Methods

Animals: C57BL/6J wild-type (WT) male and female mice (8–16 weeks old, stock no. 000664, Jackson Labs) were housed under a 12 h light / 12 h dark cycle with standard laboratory chow and water available *ad libitum*. All animal procedures were approved by the Institutional Animal Care and Use Committee of UT San Antonio and UT Southwestern Medical Center and were performed in accordance with institutional guidelines. For bulk RNA-seq studies, 32 mice were used (16 female, 16 male). A subset of male samples (n=8) was processed at UT Southwestern Medical Center. Despite correction for batch effects, these samples clustered by processing site rather than injury condition and were excluded from downstream transcriptomic analyses. For histological analyses, 16 mice were used (10 stroke (5 male, 5 female) and 6 sham (3 male, 3 female)). For axonal tracing studies, 12 mice were used (6 stroke (3 male, 3 female) and 6 sham (3 male, 3 female)).

Photothrombotic stroke: Focal ischemic lesions were induced as previously described (Labat-gest & Tomasi, 2013) via photothrombosis of vessels supplying primary motor cortex (M1). Mice were anesthetized (2-4% isoflurane /70% NO_2_ / 30% O_2_) and positioned in a stereotaxic frame. After shaving and disinfecting the scalp, a midline incision was made, and the skull overlying M1 was exposed and cleared of periosteum. Rose Bengal (40 mg/kg, intraperitoneal, Millipore Sigma, 330000-5G) was administered, and a 45 mW laser (Sapphire 561-FP OEM Laser System, 120 mW; Coherent, Santa Clara, CA, USA) was directed to stereotaxic coordinates (-1.7 ML, 0.0 AP) corresponding to M1 for 15 minutes to induce endothelial injury and local thrombus formation. At 24 h post-stroke, mice were screened for incomplete Rose Bengal delivery, and one mouse was excluded due to a subcutaneous injection, as indicated by pink discoloration of the ventral skin. Sham surgeries were performed identically to stroke procedures without the use of the laser.

Infarct volume quantification: All MRI’s were obtained by the Research Imaging Institute of UT Health San Antonio. Infarct volumes were quantified from rapid 12T MRI acquired 24 h after focal cortical stroke induction. Mice were anesthetized, and contiguous 1mm coronal slices spanning the whole brain were obtained. Hyperintense lesions were manually quantified from T2 weighted images with direct tracing and automated volume quantification using ImageJ. Stroke volume for each mouse was calculated as the sum lesion of area on each slice multiplied by slice thickness and final volumes are reported in cubic millimeters.

Tissue collection and RNA extraction: Tissue for bulk RNA sequencing was collected 7 days after stroke. Mice were deeply anesthetized and transcardially perfused with ice-cold, RNase-free 1× PBS. Brains were rapidly removed and placed on a chilled RNase-free Petri dish, and 2-mm biopsy punches were obtained from both PLC and CLC. The CLC punch was centered approximately 2 mm lateral to Bregma, whereas the PLC punch was positioned approximately 3 mm lateral and 1 mm rostral to Bregma. Tissue was immediately transferred into TRIzol reagent (Thermo Fisher Scientific) and homogenized by sequential passage through 18- and 27-gauge needles until no visible fragments remained, followed by RNA extraction as previously described (Egge et al., 2021). Briefly, total RNA was freeze/thawed/vortexed three times, followed by precipitation with chloroform and isopropanol precipitation. RNA pellets were washed twice with 75% ethanol, air-dried, and resuspended in 50 uL molecular biology grade water. A nanodrop was used to measure RNA concentration, 260/280 ratios, and 260/230 ratios. Quality control and paired-end 150bp sequencing was performed by Novogene. Only samples with RNA integrity number (RIN) ≥4, OD260/280 ≥2, OD260/230 ≥2, and no degradation or contamination were used for library preparation.

RNA-seq analysis: Libraries were sequenced as paired-end 150 bp reads on an Illumina platform by Novogene (Sacramento, CA). Reads were aligned to the mouse reference genome GRCm38, and sex specific genes were removed (i.e., Xist, Ddx3y, Eif2s3y, Kdm5d, Uty, Gm29650, Sry, Usp9y, Eif2s3x). Counts were batch corrected to account for different experimental cohorts using Combat-seq (Zhang et al., 2020). After batch correction, we performed hierarchical clustering, and samples that were segregated by processing date rather than injury (stroke vs. sham) were excluded from further analysis. 12 biological replicates (8 females, 4 males) were analyzed per condition (stroke, sham) at 7 days post-stroke. Gene-level counts were used for differential expression analysis with the DESeq2 R package, applying false discovery rate correction. Genes with ≥0.3 |log(fold change)| and FDR ≤ 0.1 were considered significantly regulated. Sequencing data are available in NCBI GEO under accession (submitted, in progress).

Weighted gene coexpression network analysis (WGCNA): WGCNA was performed using the WGCNA package (Langfelder & Horvath, 2008, 2012). Batch-corrected, variance-stabilized expression values were used as input after removing lowly expressed (genes with less than 15 counts in more than 75% of samples) and low-variance genes (variance threshold <0.01) to reduce noise. A signed coexpression network was constructed using biweight midcorrelation, soft-thresholding power β = 12, minimum module size 50, merge cut height 0.25, maximum block size 15,000, and a fixed random seed, yielding color-coded gene modules and corresponding module eigengenes. The resulting adjacency matrix was transformed into a topological overlap matrix (TOM), and genes were clustered by hierarchical clustering of TOM-based dissimilarity. Gene modules were defined by dynamic tree cutting, and closely related modules were merged based on module eigengene similarity. Module eigengenes were then correlated with experimental traits (e.g., injury status, sex, and cortical region) to identify stroke- and sex-associated gene modules. Gene ontology and pathway enrichment analyses were performed on module gene lists to infer biological processes associated with each module. Top 10 hub genes per module were identified using Cytoscape (Smoot et al., 2011) plugin CytoHubba (Chin et al., 2014), where Hubba nodes were ranked by highest degree score.

Microglia Histology: At 7 days post-stroke, mice from the sham (n=6) and stroke (n=9) groups were euthanized by isoflurane overdose and transcardially perfused with phosphate-buffered saline (PBS) followed by 4% paraformaldehyde (PFA). Brains were dissected and post-fixed in 4% PFA for 24h at 4°C, then cryoprotected in 30% sucrose in PBS. Frozen tissue was sectioned at 20 µm and mounted onto slides. Sections were washed in PBST (PBS + 0.1% Tween-20), blocked in 5% BSA in PBST, and incubated with an anti-Iba1 primary (Fujifilm Wako, 019-19741; 1:500) overnight at 4°C. Following PBST washes, sections were incubated with secondary antibody (A-11008, Invitrogen/Thermo Fisher; 1:1000). Sections were washed in PBST, coverslipped, and imaged with the Axioscan (Carl Zeiss) using consistent acquisition settings across groups. Quantification was performed using FIJI (ImageJ). Images were processed using a consistent workflow that included background correction and application of a uniform intensity threshold to isolate Iba1-positive signal. Regions of interest (ROI) were manually delineated using the freehand tool to encompass the ipsilesional secondary somatosensory cortex (S2) and the contralesional motor cortex (M1). Sections were excluded if ROIs were significantly damaged. Iba1 immunoreactivity was summarized as the area fraction within each ROI. Values were averaged across sections to generate a single value per animal. Statistical analyses were performed in GraphPad Prism, and sham versus stroke groups were compared using an unpaired two-tailed t-test.

Corticospinal tract sprouting: Four weeks after stroke induction, mice received unilateral intracortical AAV injections targeting the right primary motor cortex. Anesthetized mice (1–2% isoflurane in 70% nitrous oxide / 30% oxygen) were positioned in a stereotaxic frame and continuously monitored for temperature and respiratory rate. A small burr hole was drilled over M1, +0.5 mm anterior and 1.5 mm lateral to Bregma. A pulled glass micropipette connected to a microinjector (R-480 Nanoliter Microinjection Pump, *World Precision Instruments)* was lowered to a depth of 0.5 mm from the skull surface, and 1 µL of Synaptotag4 AAV (AAVD/J-HSyn-tdTomatoGap43-EGFPSyb2, titer ∼10^13^ gc/mL) membrane-targeted tdTomato and synaptobrevin2 fused eGFP was delivered at 100nl/min. After the virus was delivered, the pipette was left in place for 1 minute, raised 0.2mm, held again for 1 minute, and removed after injection. Buprenorphine was administered postoperatively for pain management. Moist food was provided for the first 72 hours after stroke.

At 6 weeks post-stroke, mice were transcardially perfused using room temperature PBS followed by ice-cold 4% PFA. Whole brain and spinal cords were removed, submerged in 4% PFA overnight. Whole brains were examined under a fluorescent microscope to verify injection site and expression. Animals with a visibly dim injection core were excluded from further analysis (n=6). Cervical cords were processed using the SpinalTRAQ imaging, atlasing, and analysis pipeline outlined in detail previously (Poinsatte et al., 2025). Briefly, cervical cords (pyramidal decussation to T1) were cut from the whole CNS prep and volumetrically imaged using serial two-photon tomography (STPT) on a Tissueycte 1000 (TissueVision), registered into the ST-CRA template, subjected to semi-supervised pixel classification of axons and synapses using ilastik (Berg et al., 2019), and quantified via custom R scripts for level and region-based analyses (doi:10.5281/zenodo.14750099).

## Results

### PLC and CLC transcriptomes after stroke

We performed photothrombotic stroke or sham surgery in male (n=8, 4 stroke, 4 sham) and female (n=16, 8 stroke, 8 sham) adult C57Bl6/J mice and measured T2-weighted MRI lesion volumes from a subset of mice (n=7) at 24 hours after stroke (Figure 1A). Stroke volumes ranged from 24.5 mm^3^ to 39.7 mm^3^ with no mortalities. Strokes were confined to the ipsilesional hemisphere and affected the entire primary motor cortex, part of the primary sensory cortex, and extended just below the corpus callosum (Figure 1B).

**Figure 1.**
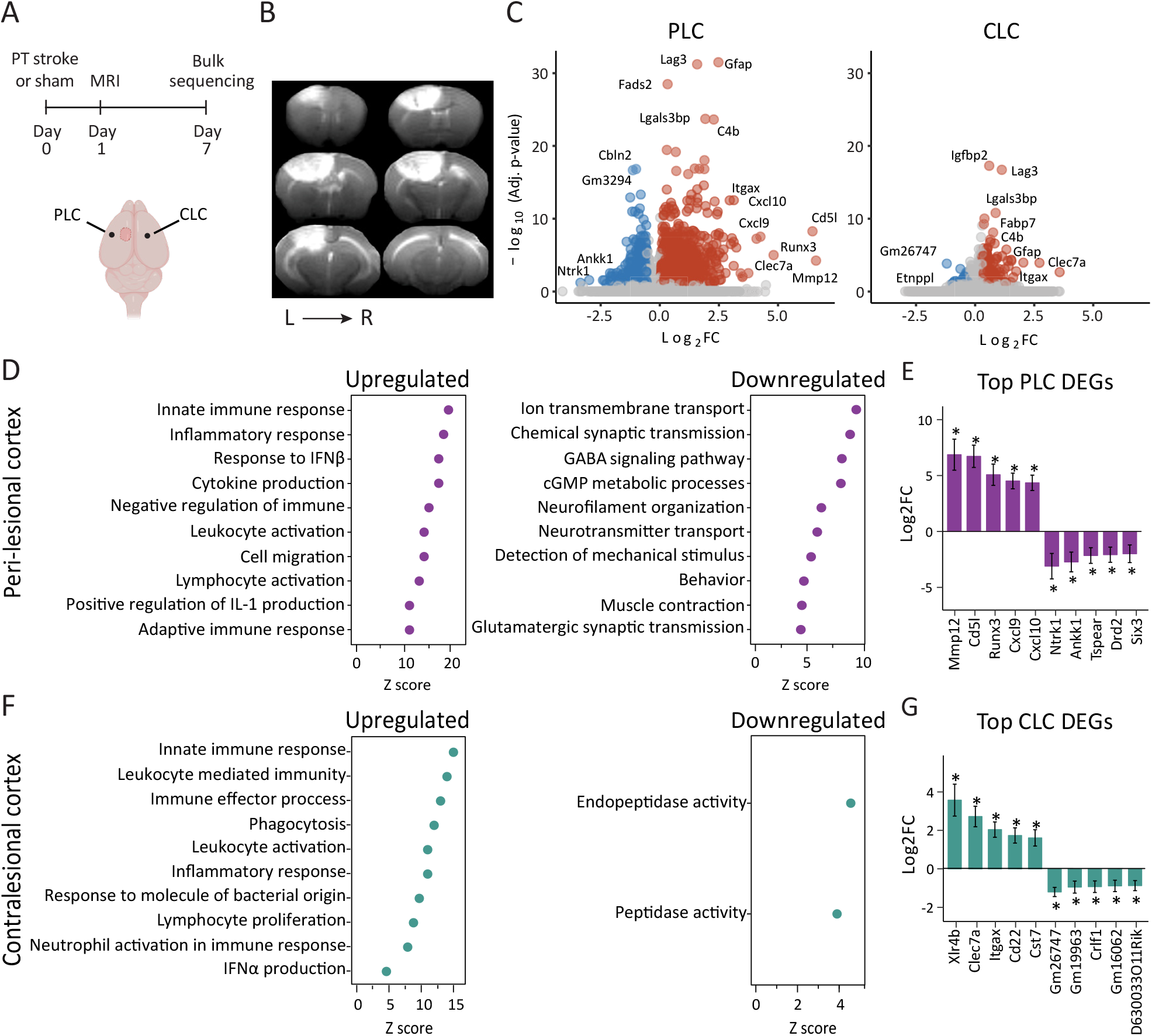
RNA-seq reveals heightened inflammatory signaling in the perilesional than the contralesional cortex 7 days post-stroke. **(A)** Experimental timeline of study and biopsy punch locations of PLC and CLC. Filled circle indicates approximate infarct location. **(B)** T2 MRI’s of representative stroke mouse with sections spanning the length of the brain rostral (top left) to caudal (bottom right). **(C)** Volcano plots for PLC and CLC (vs. respective region in sham mice, n=6/group). Upregulated DEGs (red), downregulated DEGs (blue), and nonsignificant DEGs (grey) are based on cutoff criteria of Log_2_FC ≥ 0.3 and -log_10_(FDR) < 0.1. **(D)** GO enrichment analysis of the PLC up-regulated DEGs (left) and down-regulated DEGs (right). **(E)** Bar chart of Top 5 upregulated and downregulated PLC DEGs. **(F)** GO enrichment analysis of the CLC upregulated DEGs (left) and downregulated DEGs (right). **(G)** Bar chart of Top 5 upregulated and downregulated CLC DEGs.

Region-specific transcriptomic profiles of PLC and CLC were obtained one week following primary motor cortical stroke (Figure 1). Differential expression was determined by comparing the gene expression of each site with the same region from sham-operated brains (log2FC ≥ 0.3, FDR < 0.1). Bulk RNA-sequencing of 48 samples identified 1,891 differentially expressed genes (DEGs) in the PLC (1,365 upregulated, 526 downregulated) and 274 DEGs in the CLC (245 upregulated, 29 downregulated) (Figure 1C). The PLC highly upregulated genes that align with an acute, localized response to injury, including chemokines (*Ccl2, Ccl3, Cxcl9, Cxcl10*), innate and adaptive immune system signals (*Clec7a, Fcgr4, Cd300ld, Cd300lb, Siglec1, Lilrb4a, Cd22, Ly9, Itgax, Hcar2, Fgr*), and interferon stimulated genes (*Ifi204, Ifi206, Ifi207, Ifi209, Ifi27l2a*). These genes were enriched for GO Biological Processes (GOBP) such as response to IFN-β, positive regulation of IL-1 production, cytokine production, and inflammatory response (Figure 1D, left). In contrast, downregulated genes in the PLC were linked to processes for neuronal activity and signaling, including neurofilament cytoskeletal organization, chemical synaptic transmission, and neurotransmitter transport (Figure 1D, right). Despite the remote position of the CLC relative to the lesion, highly upregulated genes also included chemokines (*Ccl3, Ccl6, Ccl9, Cxcl16*), antiviral response proteins (*Irf7, Mx1, Usp18*), and genes associated with glial reactivity and neuroinflammation (*Clec7a, Tlr2, Lag3, Itgax, Gpr84, Cd44)*. These genes were enriched for neuroinflammatory pathways such as IFN-α production, leukocyte-mediated immunity, and innate immune responses. The small set of downregulated genes in CLC (29) was enriched for GO terms related to endopeptidase and peptidase activity (Figure 1F).

We next examined the overlapping and unique DEGs for the PLC and CLC. The PLC uniquely expressed 1701 genes (1184 upregulated, 517 downregulated) that were enriched for inflammatory processes such as regulation of the innate immune response, positive regulation of cytokine production, and regulation of leukocyte activation (Figure 2A, Supplementary Figure 1). Top differentially expressed genes unique to the PLC included *Mmp12, Cd5l, Runx3, Cxcl9*, and *Cxcl10*, and *Hcar2*. Surprisingly, the CLC uniquely expressed only 84 genes (64 upregulated, 20 downregulated). Top unique upregulated genes have been linked to chromatin remodeling (*Xlr4b*) *(de la Torre-Ubieta & Bonni, 2011)*, neuronal survival (Sh3rf2) (Wilhelm et al., 2012), and synaptic integrity (Fat2) (X. Wang et al., 2025). Unique downregulated genes included Entppl, Hif3a, and Cbln4, which are involved in reactive astrocyte signaling (Tsujioka & Yamashita, 2023), hypoxia responses (Mitroshina et al., 2021), and synaptic function (Qiu et al., 2022). However, at the pathway level, GOBP enrichment of CLC-unique genes yielded broad categories, limiting interpretability (Figure 2A, Supplementary Figure 1).

**Figure 2.**
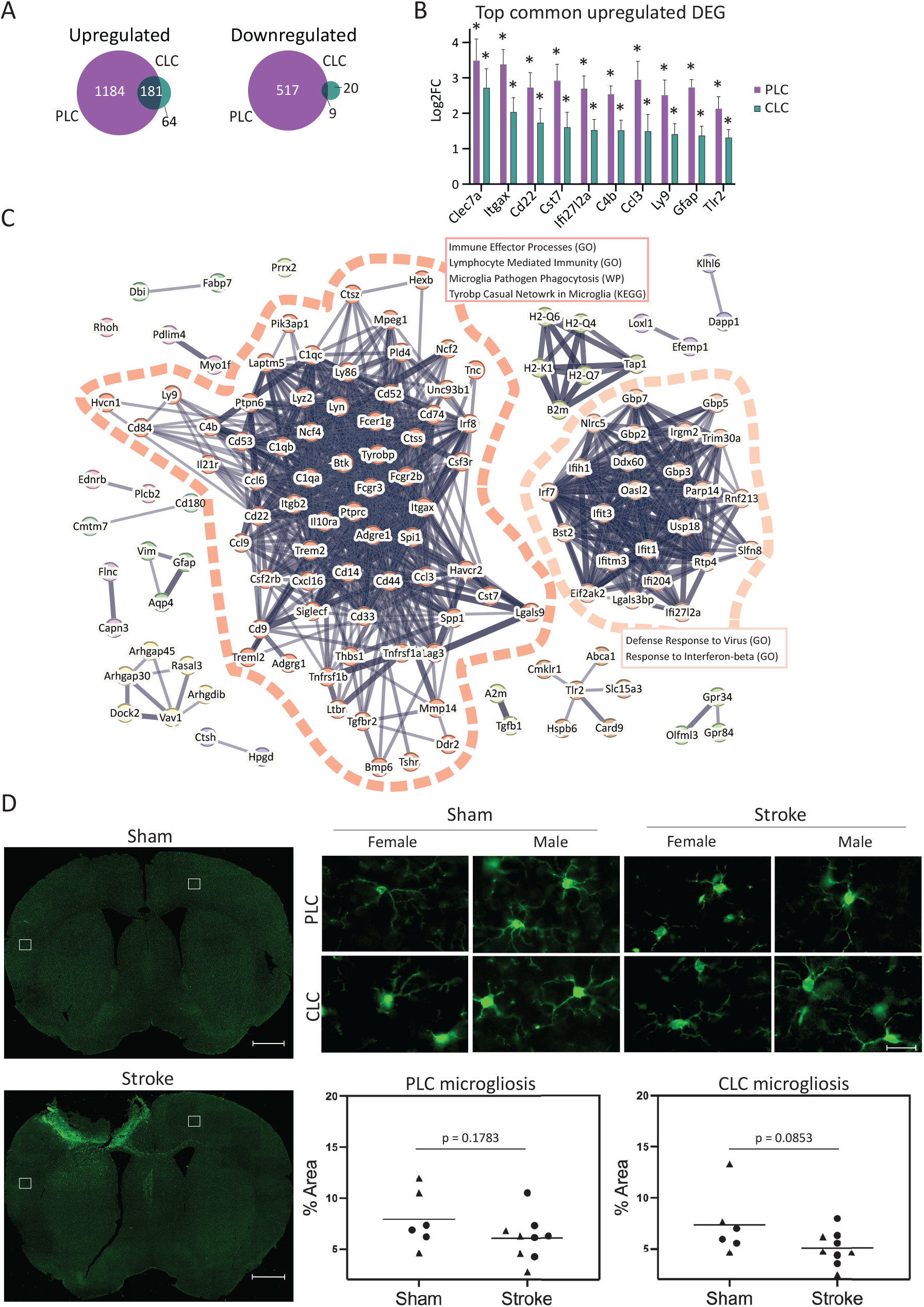
Common inflammatory signaling pathways suggest shared microglial response. **(A)** Venn diagrams comparing PLC (purple) and CLC (green) DEGs **(B)** Top shared upregulated DEGs in both PLC (purple) and CLC (green). Asterisks indicate genes that meet DEG criteria **(C)** STRING network analysis of common upregulated DEGs and pathway enrichment of largest clusters. **(D)** Representative coronal brain sections stained for IBA1 at 7 days post-stroke, shown for sham and stroke conditions in female and male mice. Quantification shows percent IBA1+ area within PLC and CLC regions of interest (ROI). Males are circles and females are triangles. Whole brain scale bar = 200 µm and zoom scale bar = 20 µm. Statistical comparisons were performed using unpaired *t*-tests (PLC: *p* = 0.1783; CLC: *p* = 0.0853)

We identified a substantial set of common DEGs across regions, indicating that CLC converges on a shared transcriptomic signature at one-week post-stroke despite its remote anatomical location. Among 181 shared upregulated DEGs, top transcripts, including *Clec7a, Itgax, Lag3, Lgals3bp, Gfap*, and *C4b*, were similarly increased in both regions (Figure 2B) and enriched for GO terms related to inflammatory activation, including IFN-α signaling, adaptive immune response, and innate immune response. Because many of these genes are markers for microglial reactivity, we next tested whether the shared signature aligned with established reactive microglial programs (Barclay et al., 2024). Shared DEGs associated with disease-associated microglial (DAM) signatures from multiple settings (Supplemental Figure 2). Consistent with this finding, STRING protein–protein interaction (PPI) analysis (Szklarczyk et al., 2019) of the common upregulated DEGs identified a large, densely connected network of reactive microglial genes that were enriched for pathways including immune effector signaling, microglial phagocytosis, and Tyrosine Protein Tyrosine Kinase Binding Protein (TYROBP)-causal network (Figure 2C). A second prominent cluster was comprised of interferon response genes (*Irf7, Ifih1, Oas2, Usp18*) that linked to anti-viral response pathways, which may arise downstream of microglial activation (Bourne et al., 2025; Depp et al., 2025; Planas, 2024). Additional peripheral networks were observed, including an astrocyte-associated group containing *Gfap, Aqp4*, and *Vim*.

Given that the shared PLC–CLC upregulated signature was largely driven by microglial-associated genes, we performed histological assessment of microglia in the PLC and CLC at 7 days post-stroke. We assessed microglia morphology by measuring the percent surface area occupied by thresholded Iba1 signal within each region. Iba1^+^ area fraction did not increase and, instead, trended lower (Figure 2D), suggesting that this microglial transcriptional program may reflect early microglial state transitions that precede morphological changes. Together, these findings indicate that reactive microglial phenotypes emerge not only as a local response to ischemic injury but also in the contralesional cortex, a remote region engaged in post-stroke plasticity.

### Transcriptional correlates of infarct volume

We performed gene-wise correlations between cortical gene expression and infarct volume to identify whether the observed transcriptional trends were associated with lesion severity. Classic injury-associated inflammatory markers, including *GFAP, IBA1*, and *IL6*, did not correlate with infarct volume in either region. While prior reports link these markers to stroke severity, they may not be sensitive enough for the small range of infarct volumes in our study (Akhoundzadeh & Shafia, 2021; Lee et al., 2025) (Supplemental Figure 3A-C). In contrast, both the PLC and CLC contained numerous transcripts whose expression was strongly associated with stroke volume (R^2^ ≥ 0.9). Many of the most highly correlated genes showed negative correlations. In the PLC, these included *Nek6* (R^2^ = 0.98, p < 0.001), a serine/threonine kinase reported to exacerbate injury by promoting astrogliosis (Yu et al., 2022), and *Fstl4* (R^2^ = 0.92), which has been implicated as a negative regulator of BDNF signaling (Suzuki et al., 2018) (Supplemental Figure 3D,E). In CLC, strongly negatively correlated genes included the deSUMOylating enzyme *SENP6* (R^2^ = 0.99), which has been linked to ischemia-induced neuronal apoptosis (Xia et al., 2021) and *Tspoap1* (R^2^ = 0.98), which has not, to our knowledge, been directly associated with stroke but participates in the regulation of neurotransmitter release (Mencacci et al., 2021) (Supplemental Figure 3F, G). Surprisingly, only one gene, *Acly*, encoding ATP-citrate lyase, a key enzyme for acetyl-CoA synthesis, was shared between regions. Acly was negatively correlated with infarct volume, consistent with prior work indicating a neuroprotective role (Y. Wang et al., 2025) (Supplemental Figure 3H). Intriguingly, the microglial activation genes identified by differential expression analyses were not among the significantly correlated genes and were not, in general, correlated with infarct volume (Supplemental Figure 3I-L).

### Hierarchical clustering of PLC and CLC transcriptomes

To determine whether gene expression signatures were consistent across animals, we performed hierarchical clustering of our 48 stroke and sham samples, which provided sufficient depth to resolve inter-animal variability and identify potential transcriptional subgroups. Hierarchical clustering of the top 50 up-regulated DEGs in both regions revealed clear separation of stroke and sham samples (Figure 3). In the PLC, stroke samples segregated into two groups. Group 1 was defined by the upregulation of genes associated with a phagocytic microglial phenotype (*Mmp12, Lgals3, Gpnmb, Clec12a*), pro-inflammatory or disease-associated microglial phenotypes (*Itgax, Cst7, Ifi27l2a, Clec7a, and C3*), and chemokines and cytokines (*Ccl3, Ccl4, Ccl5, Ccl12, Cxcl9, Cxcl10, Il1rn, and Tnfrsf26*) (Figure 3A). Group 2 was characterized by uniformly lower expression of all top 50 upregulated DEGs, indicating differences in the strength of the transcriptional response to stroke. This variation was not driven by sex or lesion severity, as Groups 1 and 2 contained both sexes and a range of infarct volumes. Hierarchical clustering of the top 50 upregulated DEGs in the CLC primarily also identified three stroke subgroups that corresponded to the PLC transcriptional subgroups (Figure 3B). Specifically, the same animals that were classified as transcriptionally distinct animals within CLC subgroup were similarly segregated in PLC subgroups. Like the PLC, CLC subgroups were distinguished by a higher expression of genes associated with reactive microglial phenotypes (*Treml2, Gpr84, Clec7a, Lgals3bp, Cst7, Lag3, Lyz3, Ifi27l2a*) (Bouchard et al., 2007; Matson et al., 2022; Zheng et al., 2016) (Figure 3B). Thus, animals distinguished by stronger microglia-associated gene expression in the PLC showed a corresponding elevation of this signature in the CLC.

**Figure 3.**
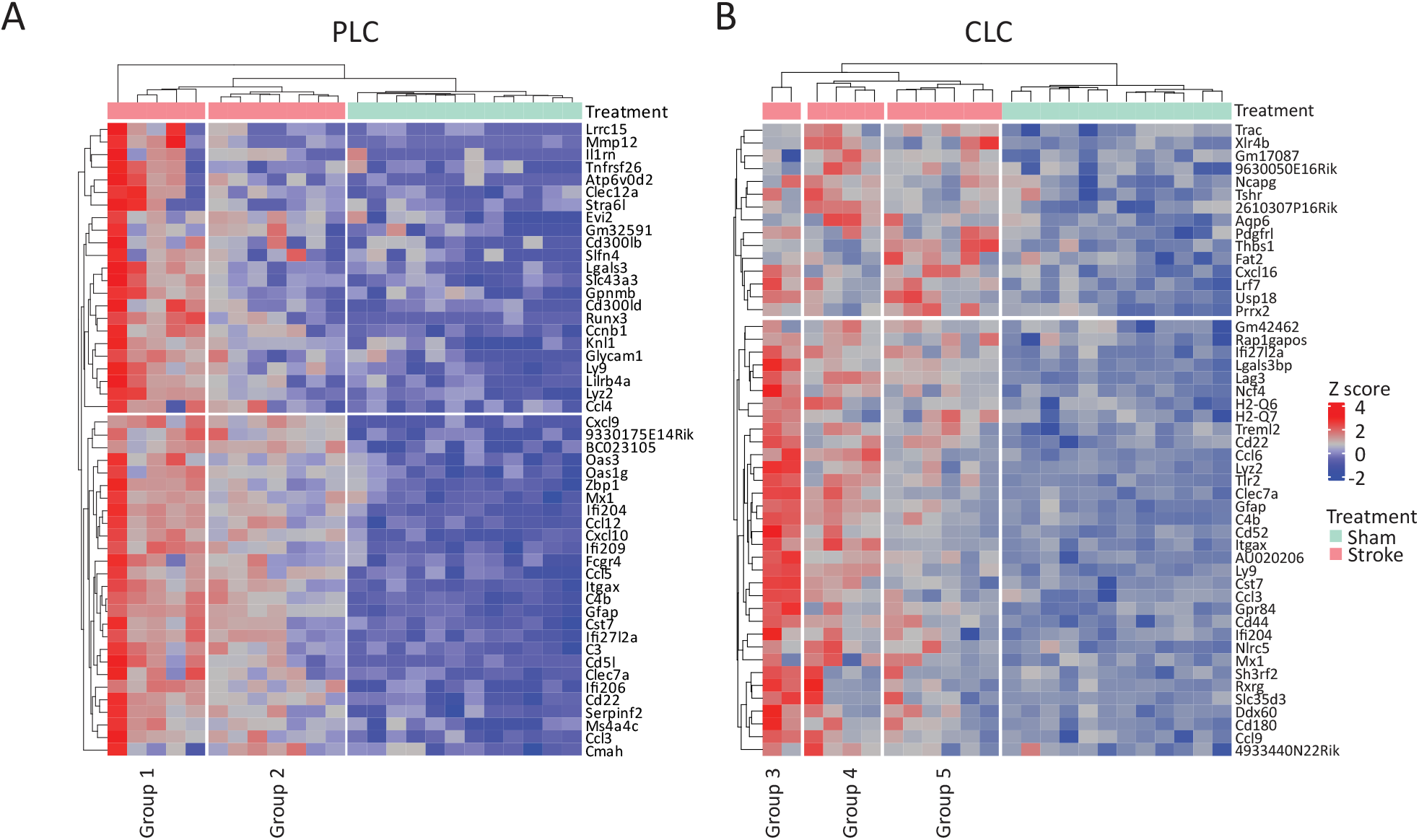
Sample-level transcriptomic analyses of PLC and contralesional cortical stroke responses. **(A-B)** Heatmap showing z-scored expression of top 50 upregulated DEGs across condition for **(A)** PLC and **(B)** CLC across samples.

### Cortical transcriptional responses to stroke in females and males

Our analyses thus far did not reveal overt sex-dependent trends. However, sex effects may be difficult to detect in pooled analyses. We therefore repeated differential expression and pathway analyses separately in male and female mice to more thoroughly evaluate sex-specific transcriptional responses. There were no differences in the lesion volumes of the subgroups measured in male vs female mice (Fig 4A). The PLC exhibited a strong transcriptional response in both females (1597 DEGs) and males (1889 DEGs) (Figure 4B). These genes were mostly upregulated and demonstrated substantial overlap (Figure 4C). GO terms for shared upregulated genes suggested a robust inflammatory response, including regulation of the innate and adaptive immune responses as well as cell proliferation and migration (Figure 4D). Highly expressed shared DEGs included Toll-like receptors (*Tlr1, Tlr2, Tlr7, Tlr9, Tlr12*), complement proteins (*C1qa, C1qb, C1qc, C1qtnf6, C3, C4b)*, cytokine and chemokines (*Ccl2, Cxcl10, Ccr5, Ccl12, Il1a*), and phagocytic effectors (*Aif1, Lgals3, Ctss, Itgam, Itgax*). GO terms for shared downregulated genes (202), which encoded synaptic proteins (*Snap25, Vamp1, Homer1, Cbln2*) and voltage-gated ion channels (*Cacna1i, Cacng2, Cacnb4, Hcn1, Scn1a, Kcna2*), included trans-synaptic signaling, GABA signaling pathway, neuronal action potential propagation, and neurofilament cytoskeleton organization (Figure 4D).

**Figure 4.**
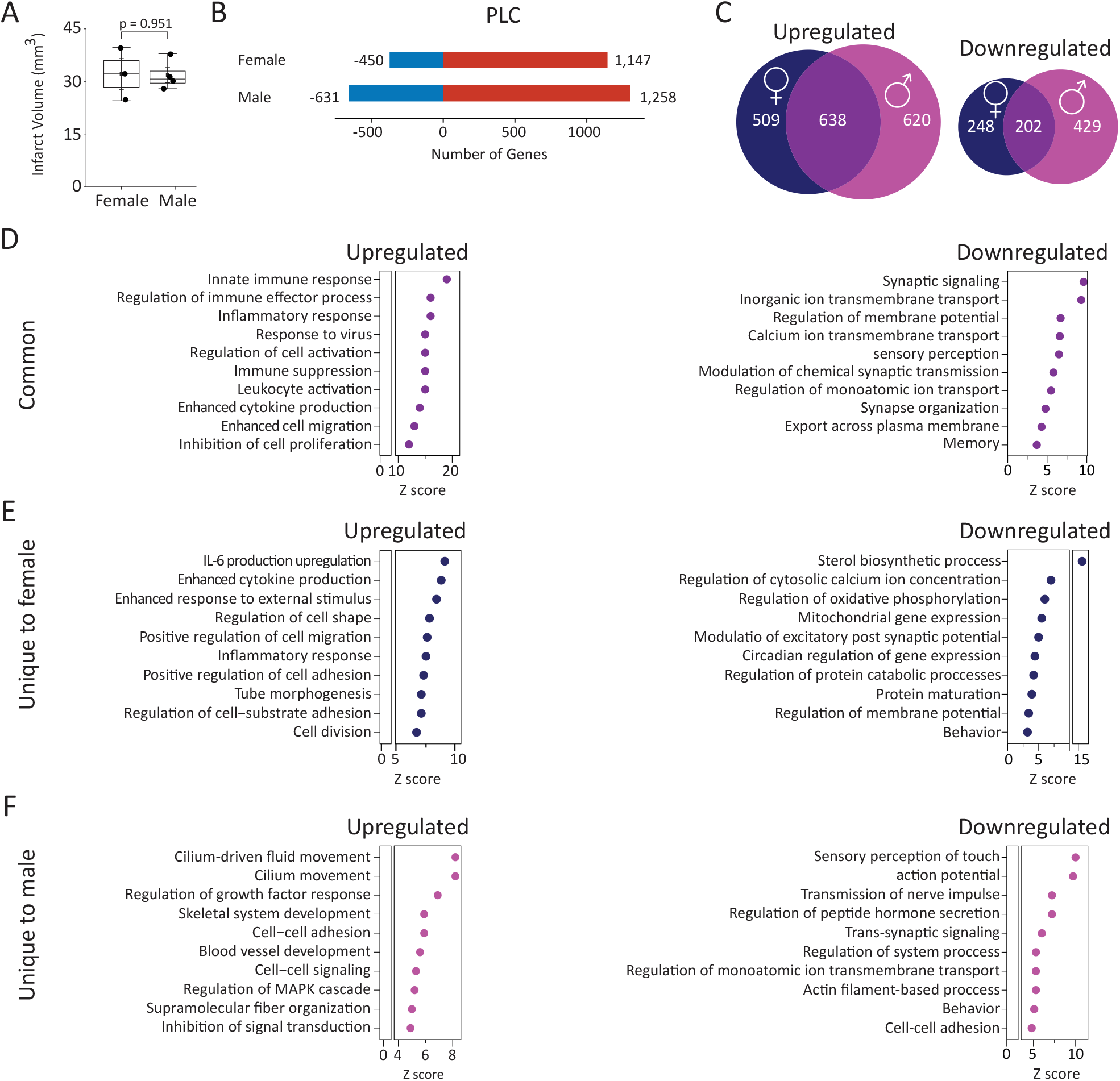
Sex-specific transcriptional responses in the PLC 7 days post-stroke. **(A)** Quantification of infarct volume 24 hrs post-stroke separated by sex (p=0.951, unpaired t-test). **(B)** PLC DEGs separated by sex, showing upregulated (red) and downregulated (blue) genes (expression relative to the PLC region in sham brains). **(C)** Venn diagrams comparing PLC DEGs across sexes, **(D-F)** GOBP enrichment for PLC DEGs that are **(D)** common among sexes, **(E)** unique to female, and **(F)** unique to male.

In the PLC, females uniquely upregulated genes encoding pattern recognition receptors (*TLR3, TLR4, TLR8, Nlrp3, Nlrp1, Rig-I, Lgp2, Aim2*) (Amarante-Mendes et al., 2018) and interferon-stimulated genes (*Ifit1, Ifitm3, Rsad2, Usp18*), suggesting increased inflammatory signaling. GO analysis of these genes indicated a positive regulation of IFN-β, IL-1, and IL-6 production (Figure 4E). Conversely, male-specific upregulated DEGs encompassed more diverse cellular processes such as synaptic signaling, astrocyte activation, and angiogenesis (Figure 4F). Downregulated genes unique to females (46) and males (227) were enriched for both neuronal signaling and plasticity. However, female-specific GO terms, including regulation of membrane potential, GABA signaling, long-term memory, and post-synapse organization, were distinct from those in males, including action potential, trans-synaptic signaling, and axon development (Figure 4F). These subtle differences may suggest a sex-dependent suppression of neuronal repair processes in the PLC.

The CLC exhibited a comparable transcriptional response in females and males, with 114 DEGs in females and 171 in males (Figure 5A). Across sexes, 43 genes were commonly upregulated, and these converged on immune-signaling pathways, including IL-6 production, leukocyte activation, and inflammatory response (Figure 5B, C). Interestingly, many of these shared transcripts have been linked to microglial activation, including *Ctss, Lag3, Ccl6, Irf7, Abca1, Cd33, Cd44, and Lgals3bp* (Griciuc et al., 2019; Kanno et al., 2005; Lei et al., 2025; Li et al., 2024; Shih et al., 2025). In contrast, only two genes were commonly downregulated (Cbln4 and Serpinb8), neither of which has a well-established connection to stroke.

**Figure 5.**
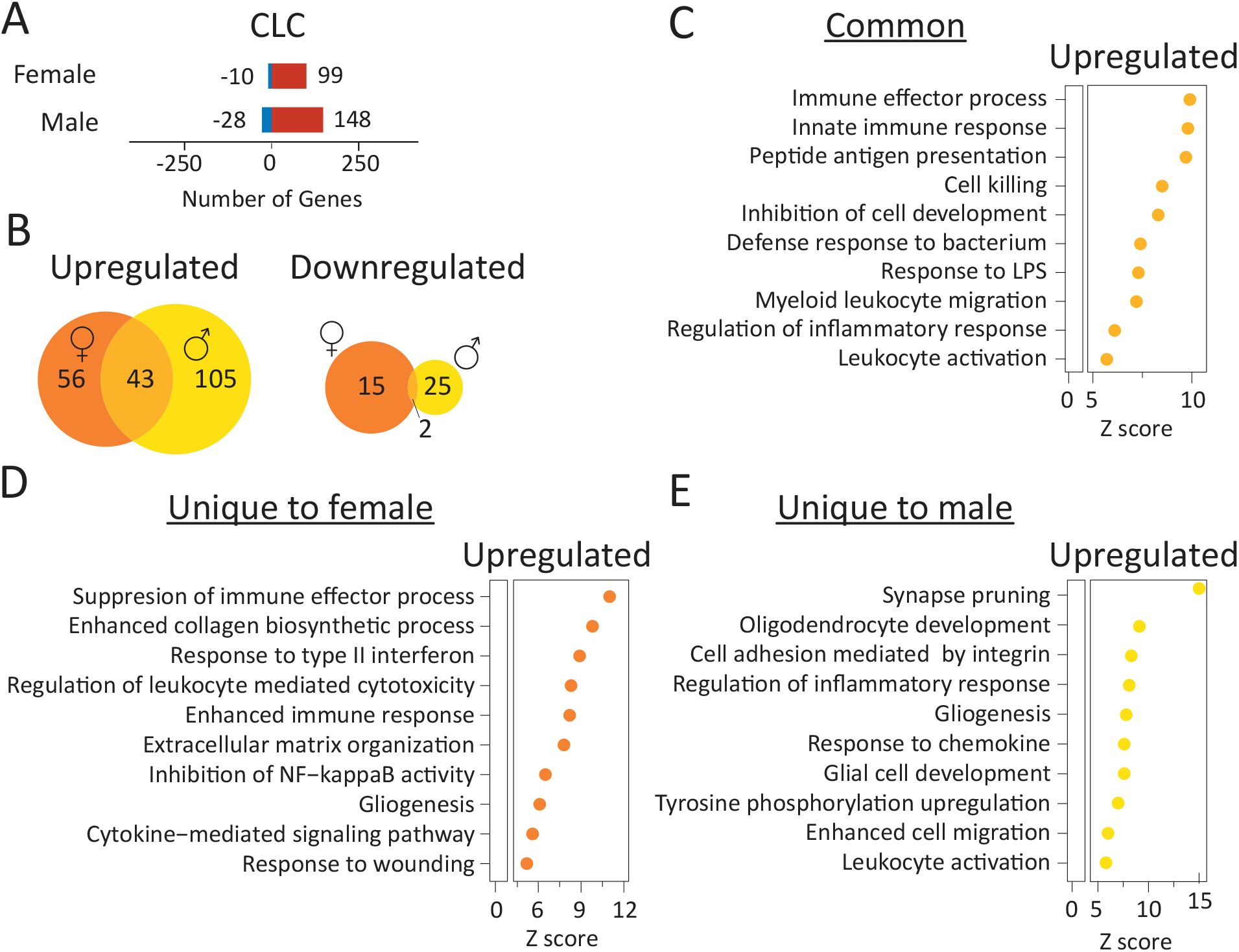
Sex-specific transcriptional responses in the CLC 7 days post-stroke. **(A)** CLC DEGs separated by sex. **(B)** Venn diagrams comparing CLC DEGs in both sexes. **(C-E)** GOBP enrichment of genes **(C)** common among sexes, **(D)** unique to female, and **(E)** unique to male

Female-specific upregulated genes (56) were enriched for pathways related to IL-1 production, response to type II interferon, regulation of the innate immune system, and vascular development (Figure 5D). Male-specific upregulated genes (105) a broader set of enriched terms, including synapse pruning, microglial activation, and inflammatory signaling (Figure 5E). Notably, gliogenesis pathways were detected in both sexes, but the contributing genes differed. In females, this signal was driven by genes such as *Gfap, Adgrg1, Sox13, Tgfb1, Vim, and Apcdd1*, whereas males expressed *C1qa, Col3a1, Csf1r, Hes5, Hexb, Kcnj10, Lyn, Nab2, Sox11, Plpp3*, and *Trem2*. Very few genes were downregulated in the CLC across both sexes and exhibited essentially no overlap.

### Coexpression network analysis of stroke- and sex-associated gene modules

To complement differential expression analysis, we examined coordinated gene expression changes via weighted gene coexpression network analysis (WGCNA). This approach identified 12 coexpression modules in the PLC and 3 in CLC (Figure 6A). In the PLC, four modules were sex-independent (Figure 6B and C). Two modules (Figure 6B) were positively correlated with stroke (“stroke-activated”) and enriched for reactive astrocyte signaling, microgliosis, and inflammatory pathways. The two negatively correlated PLC modules (Figure 6C) were enriched for neuronal pathways, including action potentials, modulation of chemical synaptic transmission, and regulation of neuronal differentiation.

**Figure 6.**
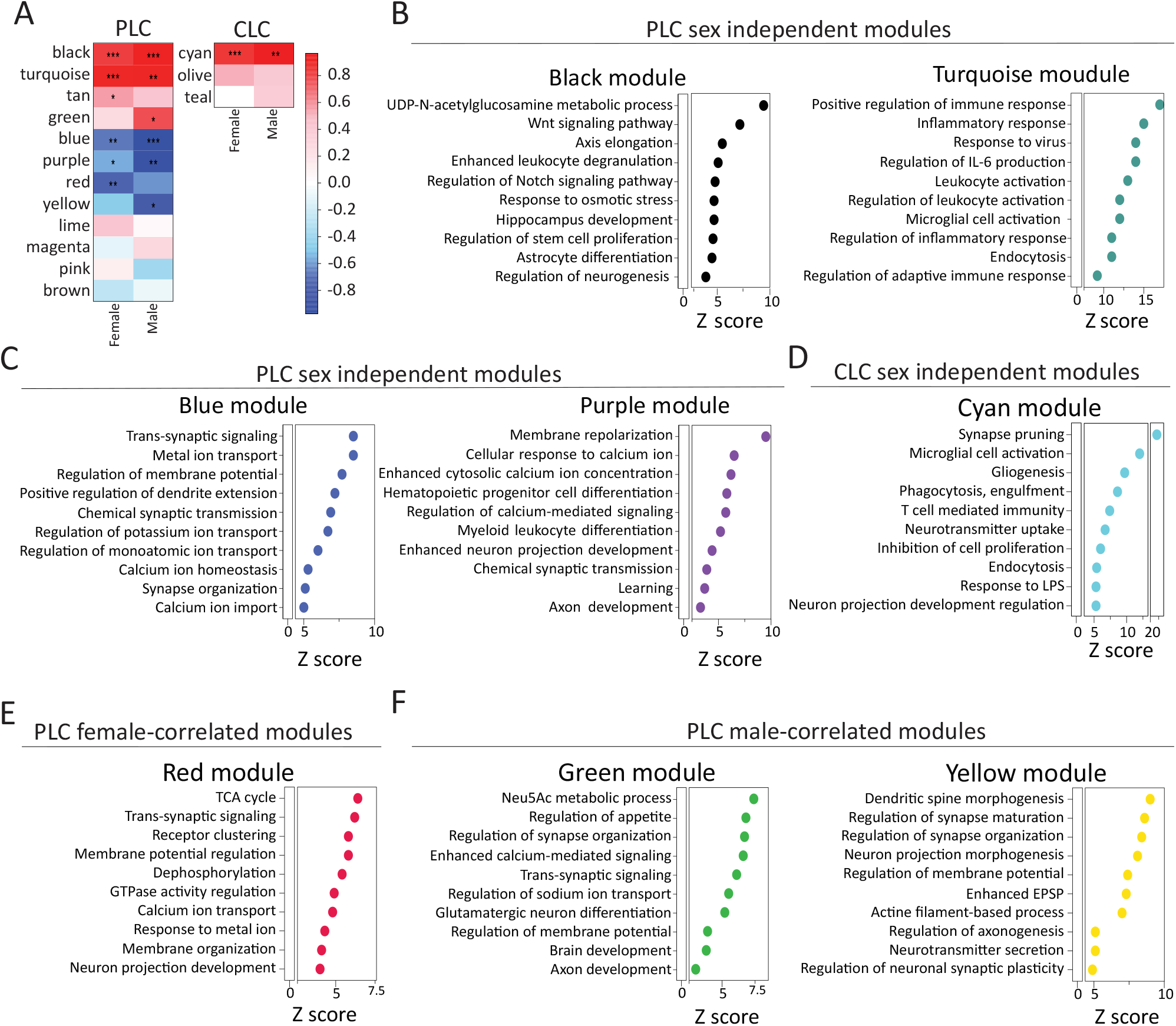
Weighted gene coexpression network analysis reveals sex-specific and regionally localized transcriptional networks following stroke. **(A)** Module-trait relationship heatmaps showing Pearson correlations between WGCNA module eigengenes and cortical region (PLC vs CLC) within stroke samples, shown separately for males and females. Color intensity indicates the strength and direction of the correlation. Asterisks indicate statistical significance (*P < 0.05, **P < 0.01, ***P < 0.001). GOBP enrichment analysis for sex-independent **(B-D)** and sex-correlated **(E-F)** modules.

Sex-dependent modules further supported suppression of neuronal signaling and plasticity in the PLC (Figure 6E and F), but through distinct gene expression. In females, the red module was negatively correlated with stroke and was enriched for pathways related to membrane potential regulation and membrane organization, and included genes involved in synaptic function (*Snap25, Gabra1, Gabrg2, Grin2c, Cnih2, Dlg1)* and protein homeostasis *(Hsph1, Rb1cc1, Vmp1, Hspa4, Ube2g1*) (Figure 6E). In males, the yellow module was negatively correlated and included transcripts linked to axon guidance cues (*Epha5, Ephb3, Sema3f, Nrp2, Unc5a, NogoR*), ion channels (*Kcnq4, Kcnb1, Trpm4*), and cellular stress response proteins (*Xrcc4, Prdx1, Nqo1, Sesn3*) (Figure 6F). Together, these sex-specific modules suggest that suppression of neuronal plasticity in the PLC may affect synaptic signaling in females and structural remodeling in males.

While PLC gene expression has consistently suggested a suppression of neuronal signaling and plasticity under stroke conditions, one male-specific positively correlated module (Figure 6F) was upregulated in the PLC and was notable for genes linked to neuronal growth and repair; this module contained 182 upregulated genes, including *Esr1*, which was upregulated in males but not females, and featured hub genes such as *GAP43, Esr1, and Calb2*. Females exhibited one positively correlated module (tan); however, its members were enriched for leptomeningeal, choroid-plexus, and extracellular matrix markers (*Cdh1, Fbln1, Foxc1*), which can sometimes indicate variation in tissue composition or sample handling (Olney et al., 2022).

In CLC, WGCNA yielded three modules, of which only one (cyan) was significantly associated with stroke (Figure 6F). This module was enriched for microglial activation, synaptic pruning, and gliogenesis. The hub genes for the cyan module (*Cd68, Gfap, Apoe, Csf1r, Fcgr3, B2m, and Cd44*) suggest microglial- and astrocyte-mediated inflammatory signaling in the CLC across both sexes. Overall, WGCNA supports findings of sex-independent inflammatory signaling and suppression of neuronal plasticity in the PLC and a glial-driven neuroinflammatory response in the CLC.

### Post-stroke CST plasticity across sexes

We next asked whether sex-dependent transcriptional programs correspond to differences in post-stroke plasticity. To examine this question, we quantified axonal sprouting of the contralesional corticospinal tract (CST), a robust, functionally relevant form of post-stroke plasticity, in both female and male mice. Using our previously developed spinal cord analysis pipeline, SpinalTRAQ (Poinsatte et al., 2025), we mapped and quantified contralesional CST synapses throughout the entire cervical spinal cord (Figure 7A). In sham animals, both sexes showed strikingly similar CST innervation patterns, with higher normalized synapse density in dorsal and intermediate laminae (laminae 5, 7–10) across C1–C8 and minimal innervation of dorsal laminae 1–2 (Figure 7B,C). Six weeks after stroke, CST sprouting and synapse formation into the denervated hemicord significantly increased in both females and males. In line with our previous findings (Poinsatte et al., 2025), both sexes showed the largest gain in synapses in lamina 5 and 7 across all cervical levels after stroke. Additionally, we found increases in lamina 4 and 9 synapses that localized to caudal cervical sections. This indicates that, despite robust differences in cortical CLC gene expression between males and females, we found no evidence for sex differences in contralesional CST synaptic plasticity at the level of the cervical spinal cord.

**Figure 7.**
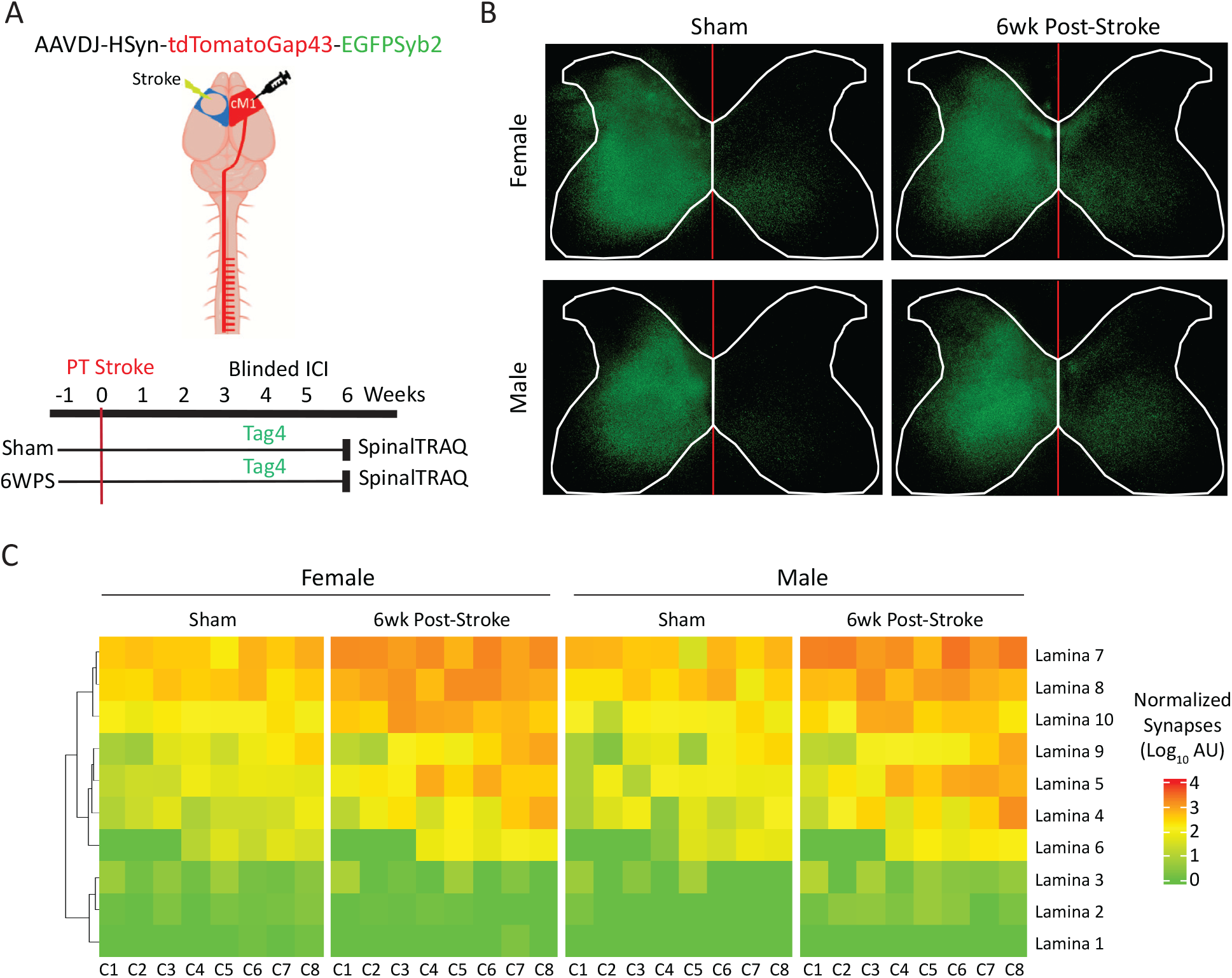
Comparison of chronic post-stroke spinal plasticity by sex. **(A)** Experimental schematic: adult C57BL/6 mice (n = 3 per group) underwent PT stroke or sham surgery followed by anterograde synaptic labelling (Synaptotag4) at 4 weeks post-stroke and SpinalTRAQ analysis at 6 weeks post-stroke. **(B)** Maximum intensity projections (MIPs) of synapse-classified pixels from the volumetric cervical spinal cord dataset of a representative animal. White outlines denote spinal cord boundaries, and the red line indicates the midline. **(C)** Heatmap of classified synaptic signal showing group-mean normalized intensity (log_10_ scale) across cervical spinal levels and lamina. Laminae are hierarchically clustered based on presynaptic terminal distributions in the contralateral hemicord.

## Discussion

Stroke induces plasticity in both lesion-adjacent and lesion-remote regions. In rodent models, one of the most well-characterized remote regions is the contralesional cortex (CLC). The CLC is truly distinct from the lesion and lacks direct damage. Nevertheless, our lab and others have found that significant post-stroke plasticity occurs in this region, similar to lesion-adjacent regions. However, it remains unclear whether the CLC engages distinct molecular pathways. In this study, we used bulk RNA sequencing at 7 days post-stroke to compare the peri- and contra-lesional transcriptomes.

Consistent with prior work, the PLC exhibited a broad induction of innate and adaptive immune pathways and suppression of neuronal genes (Choi et al., 2019; Lu et al., 2003; Ramos-Cejudo et al., 2012; Rust, 2023). While the CLC displayed a significantly more modest response, its gene expression signature indicated robust activation of a glial-derived neuroinflammatory response and, unlike the PLC, preserved neuronal gene expression. Both the PLC and CLC converged on a regionally localized transcriptional signature characterized by inflammatory and microglial activation pathways. This shared gene expression included markers for multiple microglial phenotypes, including reactive, disease-associated (DAM) microglia. These results highlight reactive microglial signaling as a prominent pathway in both regions. Next, we investigated transcriptional responses in male and female mice. Although there were no significant differences in infarct volumes, inflammatory gene expression was more robust in the PLC and CLC in female mice. Male mice uniquely upregulated genes linked to synaptic signaling, angiogenesis, and glial activation in both regions, a finding further supported by WGCNA, which revealed a unique post-stroke module enriched for synaptic plasticity pathways. Despite these differences in gene expression, we found that post-stroke plasticity was robust in both sexes and exhibited no major differences in targeting or magnitude. Overall, our data support a model in which stroke induces regionally localized inflammatory responses that may be permissive for post-stroke plasticity.

While changes in circuit activity can be detected within hours to days after stroke, large-scale structural plasticity is typically detected at later timepoints, including weeks to months after stroke (Carmichael et al., 2001; Dancause et al., 2005; Lapash Daniels et al., 2009; Poinsatte et al., 2025; Zhang et al., 2010). Accordingly, we chose to examine gene expression at day 7 post-stroke, a timepoint commonly considered an initiation phase for neurons to begin a phenotypic shift that supports later remodeling, including axonal sprouting (Li et al., 2010) . We anticipated robust induction of neuronal plasticity programs at this stage; however, most differentially expressed genes reflected non-neuronal cell types, and in the PLC we observed suppression of genes linked to neuronal plasticity-related processes. This suggests that bulk RNA sequencing is limited for detecting neuronal transcriptional responses to CNS injury, particularly when injury-driven shifts in cellular composition dilute neuronal signal. In addition, neurons experiencing a major disruption of connections may mount transcriptional changes that differ from those that sustain direct axonal injury, and these distinct responses may be masked when bulk profiling averages signals across heterogeneous neuronal states and extrinsic cues.

The CLC is unique in that it is remote from the lesion, yet it exhibits robust neuronal plasticity, including axonal sprouting, which is limited in the mature nervous system. This led us to hypothesize that the CLC would exhibit a transcriptomic program that differs from a lesion-adjacent region, where neurons are recovering from direct injury and exposed to a highly inflammatory milieu. Unexpectedly, the CLC showed a largely overlapping transcriptional response with the PLC. Although the PLC exhibited substantially greater peripheral immune activation, both regions shared transcriptional features consistent with microglial reactivity. In particular, GO term enrichment among commonly differentially expressed genes highlighted cytokine and pro-inflammatory signaling pathways, which are frequently associated with activated microglial states (Bourne et al., 2025; Tang et al., 2025). Several of the top shared DEGs were also closely linked to reactive microglial phenotypes, including DAM-associated transcripts such as Clec7a, Cst7, and Itgax (Barclay et al., 2024; Depp et al., 2025). DAM-like microglial populations have been reported in remote post-stroke regions on the ipsilesional side (Cao et al., 2021), and while their role in stroke recovery remains incompletely defined, they have been proposed to reflect a phagocytic state necessary for structural reorganization (Depp et al., 2025; Jia et al., 2023; Wan et al., 2024). We also observed marked upregulation of Ifi27l2a, a regulator associated with pro-inflammatory microglia that has been reported in PLC (Kim et al., 2023). Its prominence in CLC supports the possibility that pro-inflammatory microglial states can extend beyond the PLC to influence post-stroke remodeling. Finally, the shared upregulation of Ccl3, Tlr2, and Cd22, genes implicated in both peripheral immune and reactive microglial signaling, further suggests microglial reactivity and supports prior work establishing immune cell infiltration in remote regions after stroke (Kang et al., 2018; Pluvinage et al., 2019; Selvaraj et al., 2021; Tang et al., 2007; Torres et al., 2025).

We next investigated how the transcriptional responses to stroke varied by sex, especially since studies in female mice are extremely limited (Rust, 2023). In the PLC, both sexes showed robust induction of innate and adaptive immune pathways and downregulation of synaptic and excitability genes. However, females exhibited stronger upregulation of pattern recognition receptors and interferon-stimulated genes, with GO terms pointing to enhanced IFN-β, IL-1, and IL-6 production. Males, in contrast, showed additional enrichment for synaptic signaling, astrocyte activation, and angiogenesis. WGCNA extended this view by identifying female-specific modules with suppressed pathways for synaptic signaling and excitability, and a male-specific module enriched for neuronal growth and repair, with hub genes such as GAP43, Calb2, and Esr1. While these data may support the idea that males and females exhibit partially distinct molecular processes, more extensive investigation is needed to determine whether these differences would be useful for sex-specific interventions. It is important to note that these sex-dependent transcriptional differences did not translate into large-scale differences in structural plasticity of the contralesional corticospinal tract. Assessment of the whole cervical spinal cord with SpinalTRAQ found that both females and males exhibited comparable contralesional CST sprouting into the denervated hemicord at 6 weeks post-stroke. Baseline CST innervation patterns were also similar between sexes. Thus, the stronger inflammatory activation seen in females in both the PLC and CLC did not result in differences in our model of post-stroke plasticity. This may be due to sex-dependent differences in timelines or compensatory mechanisms. Alternatively, the sex differences we observed in the brain may influence other aspects of recovery that are not captured by contralesional CST sprouting.

Several limitations restrict the interpretation of these findings. First, transcriptomic profiling was restricted to a single subacute time point. We therefore cannot determine whether sex or region-specific differences are transient or persistent. Second, we relied on bulk RNA-seq from heterogeneous cortical tissue. Although prior work and cell-type specific markers provide insight into underlying cell types, we cannot definitively tie changes to specific microglial, astrocytic, neuronal, or peripheral immune populations. Third, our analyses are limited to mRNA and do not address protein levels or functional effects of changes in gene expression. Fourth, our measure of post-stroke plasticity focuses on one major form of plasticity, axonal sprouting, from a single pathway that contributes to stroke recovery, the uninjured CST. We did not assess other circuits or forms of plasticity that may differ by sex. Finally, we did not directly link individual transcriptional modules or candidate genes to behavioral outcomes. Future work should address these gaps by performing single-cell level assessments, such as single-nucleus or spatial transcriptomics, with parallel measurements of plasticity and behavior. Profiling across acute, subacute, and chronic phases will be critical to understand which cell-type specific pathways contribute to stroke recovery and whether these roles differ by sex, region, or time. Single-cell and spatial approaches have identified key microglial, astrocytic, neuronal, and vascular contributions after stroke, but typically have not involved a comparison of multiple regions within the same study (Bormann et al., 2024; Garcia-Bonilla et al., 2024; Han et al., 2024; Ikegami et al., 2025; Jin et al., 2023; Kim et al., 2023; Scott et al., 2024; Zheng et al., 2022). Furthermore, targeted manipulation of candidate modules is an important addition to these studies, particularly when combined with CST mapping and behavioral assays in order to differentiate correlates of recovery versus active drivers. Finally, extending these approaches to other stroke models, like those inducing post-stroke cognitive decline, would be useful for understanding shared, circuit- or cellular-level responses across stroke subtypes.

In summary, this work defines a shared cortical transcriptional state at day 7 after focal motor cortical stroke, characterized by microglial-mediated inflammation in the PLC and CLC. Broad gene expression patterns and targeted contralesional CST sprouting in females and males were similar. These findings support the idea that post-stroke inflammation is more than a unilateral response to direct injury. Rather, inflammatory programs are broadly active across regions that undergo post-stroke plasticity, where they likely play a critical role in stimulating or modulating structural reorganization. By linking unbiased cortical transcriptomics with cervical spinal cord mapping of CST plasticity, our study provides a framework for identifying molecular targets that enhance recovery, while also highlighting specific axes where future mechanistic studies should focus, such as regions, cell types, and sex. This approach also creates opportunities to identify convergent molecular signatures across spatially remote brain regions, potentially enabling early biomarkers for plasticity and repair after stroke and other CNS injury conditions.

## List of abbreviations

(PLC): Perilesional Cortex
(CLC): Contralesional Cortex
(PT): Photothrombotic
(RNA-seq): RNA sequencing
(PPI): protein–protein interaction
(DAM): Disease-associated Microglia
(EAE): experimental autoimmune encephalomyelitis
(5xFAD): 5 Familial Alzheimer’s Disease
(GOBP): Gene Onotology Biological Processes
(GEO): Gene Expression Omnibus

## Acknowledgements

LLM tools (ChatGPT) were used for basic editing and grammatical refinement. We’d like to thank Dr. Kumar Sharma and Ian Tamayo for assistance with whole-slide imaging.

## Notes

Funding: This work was supported by National Institutes of Health (NIH) awards T32AG082661 (DB), T32GM -113896 and -145432 (DB), T32NS082145-08 (DB), AHA Fellowship 23PRE1018993 (DB), GSBS Neuroscience Training Fellowship (MK), NS088555 (AMS), OT2OD032581 (MPG), and by gifts from the Haggerty Foundation (MPG) and Meier Family Foundation (MPG). P.M.D and the members of the Douglas lab were supported by the Clayton Foundation for Research, the Welch Foundation (I -2061 -20210327), the American Federation of Aging Research (AFAR 2023), the National Institutes of Health (R01AG076529, R01GM15385)

### Competing Interest Statement

The authors have declared no competing interest.

