## Supplementary figures and images for "Shared Transcriptomic Signatures in Perilesional and Contralesional Cortex"

### Supplementary Figure 1

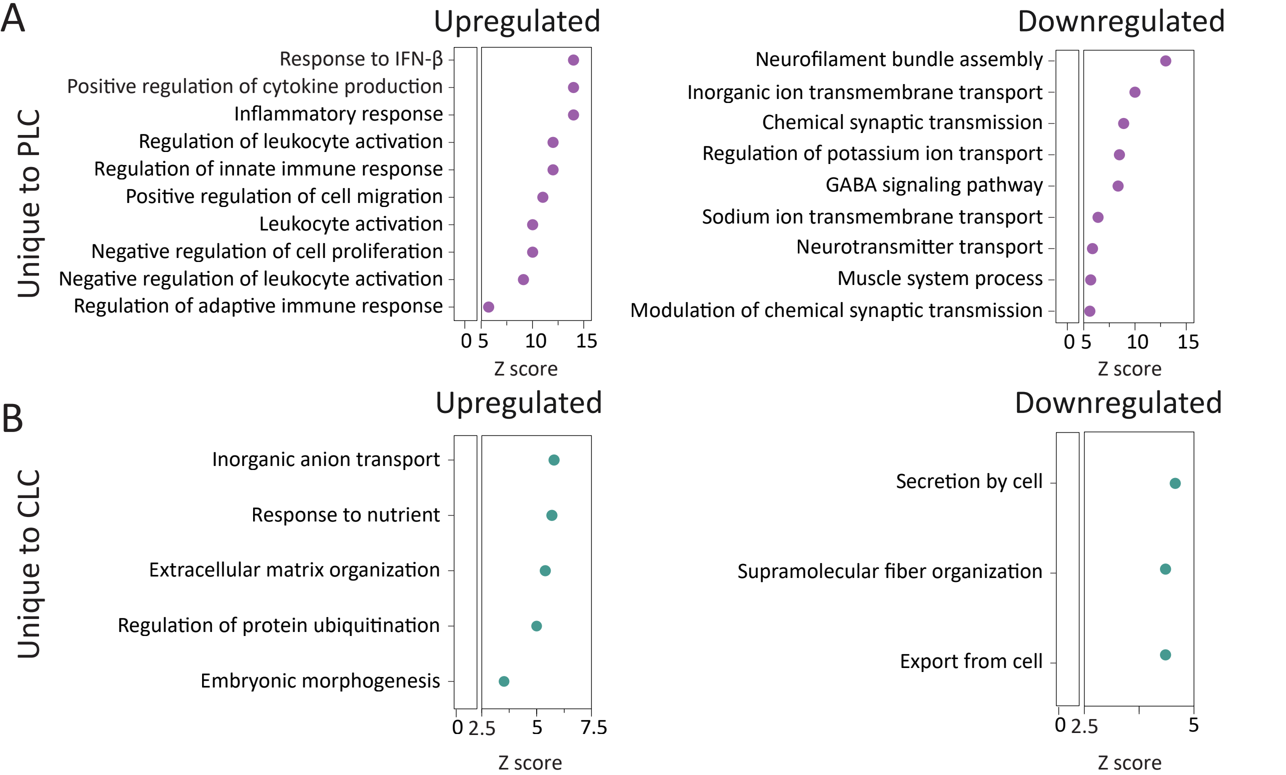

### SupplementaryFigure 2

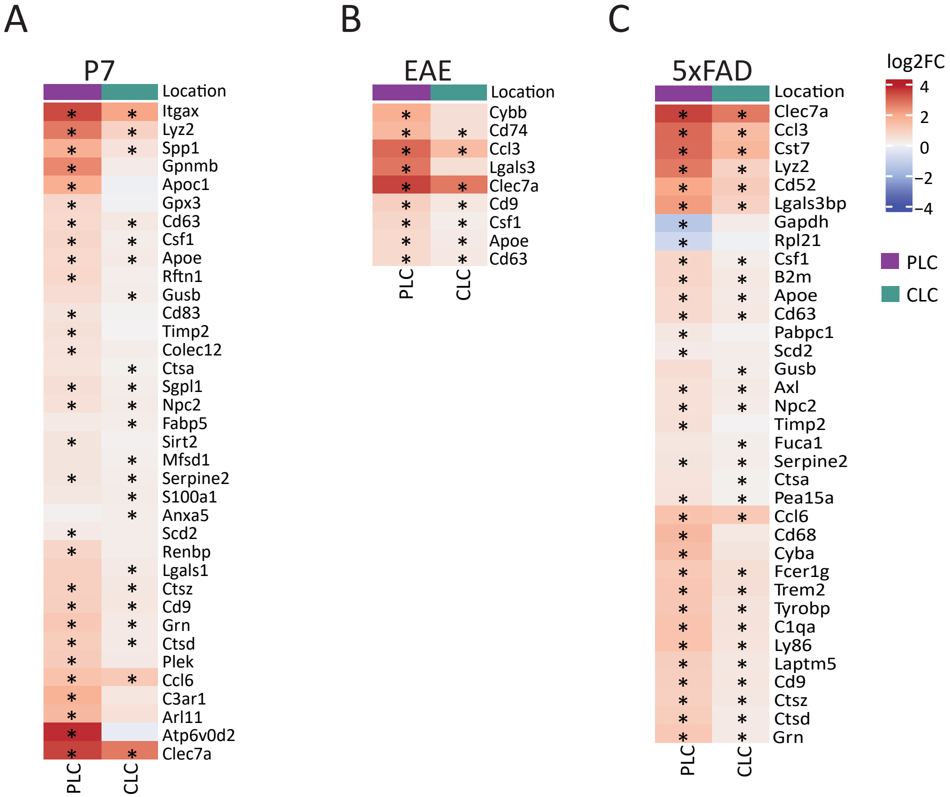

### SupplementaryFigure 3

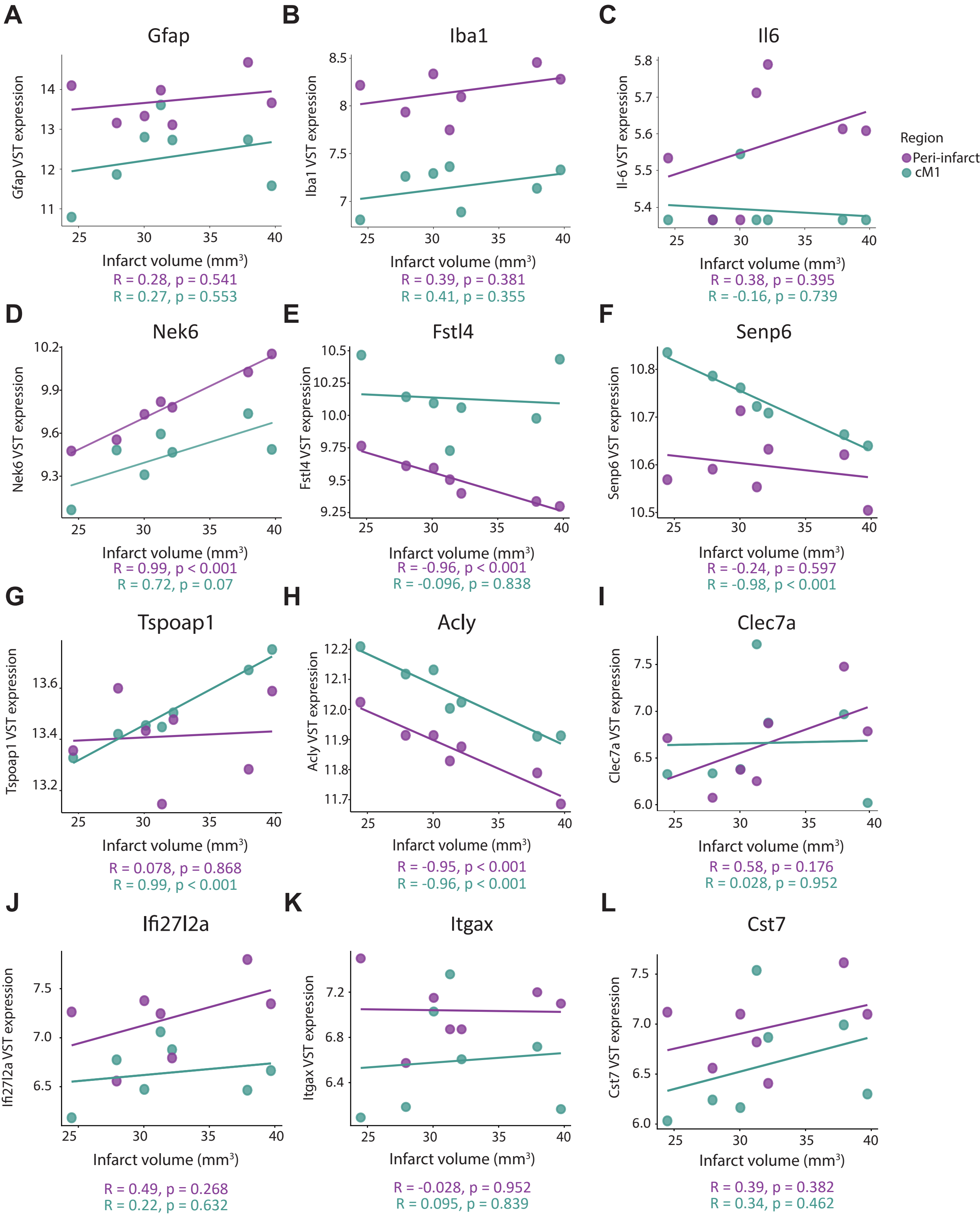
